# Fish perform like mammals and birds in inhibitory motor control tasks

**DOI:** 10.1101/188359

**Authors:** Tyrone Lucon-Xiccato, Elia Gatto, Angelo Bisazza

## Abstract

Inhibitory control is an executive function that positively predicts performance in several cognitive tasks and has been considered typical of vertebrates with large and complex nervous systems such as primates. However, evidence is growing that some fish species have evolved complex cognitive abilities in spite of their relatively small brain size. We tested whether fish might also show enhanced inhibitory control by subjecting guppies, *Poecilia reticulata*, to the motor task used to test warm-blooded vertebrates. Guppies were trained to enter a horizontal opaque cylinder to reach a food reward; then, the cylinder was replaced by a transparent one, and subjects needed to inhibit the response to pass thought the transparency to reach the food. Guppies performed correctly in 58 % of trials, a performance fully comparable to that observed in most birds and mammals. In experiment 2, we tested guppies in a task with a different type of reward, a group of conspecifics. Guppies rapidly learned to detour a transparent barrier to reach the social reward with a performance close to that of experiment 1. Our study suggests that efficient inhibitory control is shown also by fish, and its variation between-species is only partially explained by variation in brain size.

## Introduction

Inhibitory control is one the core executive functions and allows an animal to control attention and behaviour in order to override internal predispositions or resist to external lures [1, 2, 3]. One of the most studied aspects of this executive function is inhibitory motor control, which is required when one individual has to block an impulsive behaviour [4]. Inhibitory control has been shown to correlate with performance in many cognitive tasks, and it is believed to be a prerequisite for sophisticated cognitive skills. For example, performance in tasks requiring inhibition correlates with intelligence in adult humans [5], and in children it positively predicts academic achievement along with cognitive competence in later life [6, 7, 8]. In cotton-top tamarin, *Saguinus oedipus*, inhibitory control predicts problem-solving performance [9], whereas in song sparrows, *Melospiza melodia*, it predicts song repertoire size [10].

Efficient inhibitory control has often been considered a distinctive feature of humans [11] or vertebrates with large, complex nervous systems [12], as observed for other important cognitive abilities [13, 14]. This idea is mainly based on evidence that even children and non-human primates often show difficulties in solving inhibitory control tasks [15, 16, 17, 18, 19]. Empirical support has been provided recently by a comprehensive study on three dozen mammalian and avian species, demonstrating that inhibitory performance positively correlates with brain size [20]. However, most of the species tested in that study were mammals, and subsequent research has shown that other bird species perform similarly to apes despite the much smaller brain size [21].

Several complex cognitive processes and abilities believed to be distinctive of few mammalian and avian species have been recently observed in teleost fish in spite of their relatively small brain size [22, 23]. For instance, some fish species use tools, transmit cultural information, show problem solving, learn complex spatial mazes, and display episodic-like memory [22, 24, 25, 26, 27, 28]. Some of the cognitive tasks successfully solved by fish, such as reversal learning, require inhibition to some extent [29, 30]. Given the suggested link between inhibition and cognitive attainment in other tasks [5, 6, 7, 8, 9], we hypothesised that inhibitory control might be elevated in fish too, at least in those species that have evolved notable cognitive abilities. To address this hypothesis, we investigated inhibitory control in the guppy, *Poecilia reticulata*, using two motor tasks that exploit the response of animals to the presence of transparent obstacles. The guppy is an ecological generalist species characterized by a considerable behavioural flexibility that has permitted the successful invasion of many different environments in all continents outside Antarctica [31]. Guppies have been shown to be capable of complex behaviours and enhanced cognitive abilities [28, 32, 33].

In experiment 1, we tested guppies using the cylinder task, which has been widely adopted to study inhibitory control in mammals and birds [10, 20, 34]. We followed the procedure adopted by MacLean and colleagues to compare 32 different species [20]. Guppies were initially trained to enter a horizontal opaque cylinder to reach a food reward; in the following test trials, guppies were presented with a transparent cylinder and had to enter the cylinder from the open lateral sides overcoming the tendency to swim directly toward the visible target. MacLean and colleagues [20] performed short experiments (10 test trials), but some studies have suggested that animals can increase their performance over trials in similar inhibitory tasks [21, 35]. To study the occurrence of learning in guppies, we lengthened the duration of the experiments up to 50 trials (5 trials per day). In the study by MacLean and colleagues [20], the cylinder task was considered a self-control task. However, some authors have argued that self-control is required when an individual has to resist to the temptation to obtain an objectively less valuable target in order to obtain a more valuable target after a temporal delay [3, 36]; following these authors, we considered the cylinder task as a measure of inhibitory motor control.

Recent studies have recommended the use of multiple tests for assessing the cognitive abilities of a species [37, 38, 39]. This appears to be particularly important in the case of inhibition because the performance of the different species may vary according to the relative value of the reward [40]. For example, the different food intake requirements of warm- and cold-blooded species might affect the performance in tasks using food as an attractor. Thus, in experiment 2, we tested guppies in an inhibitory motor control task that uses a social stimulus, the barrier task. We based our procedure on the test adopted to study spatial abilities and lateralisation in guppies and in other fish species [41, 42]. In a series of 25 test trials, guppies were inserted in a novel tank and had to detour a C-shaped transparent barrier to reach a shoal of conspecifics.

## Results

### Experiment 1: cylinder task

In the training phase with the opaque cylinder, guppies reached the learning criterion of 4 out of 5 daily correct trials after 17.5 ± 8.25 trials (mean ± SD). During the entire test phase, guppies performed 58.40 ± 11.07 % trials in which they attempted to retrieve food from the side of the transparent cylinder rather than through the transparency (correct trials). The proportion of correct trials in the test phase was significantly lower compared to that in the last day of the training phase (paired-sample *t* test: *t*_9_ = 7.589, *P* < 0.0001). The performance of the individual fish ranged between 38-72 % correct trials. The likelihood of correct trials did not significantly change across trials (GLMM: *χ*^2^_1_ = 1.350, *P* = 0.245; Fig. 1a), but the time to enter the cylinder significantly decreased across trials (LMM: *χ*^2^_1_ = 8.668, *P* = 0.003; Fig. 1b). Considering only the initial 10 trials, as in the study by MacLean and colleagues [20], guppies performed 53.00 ± 29.83 correct trials (Fig. 2).

**Fig. 1.**
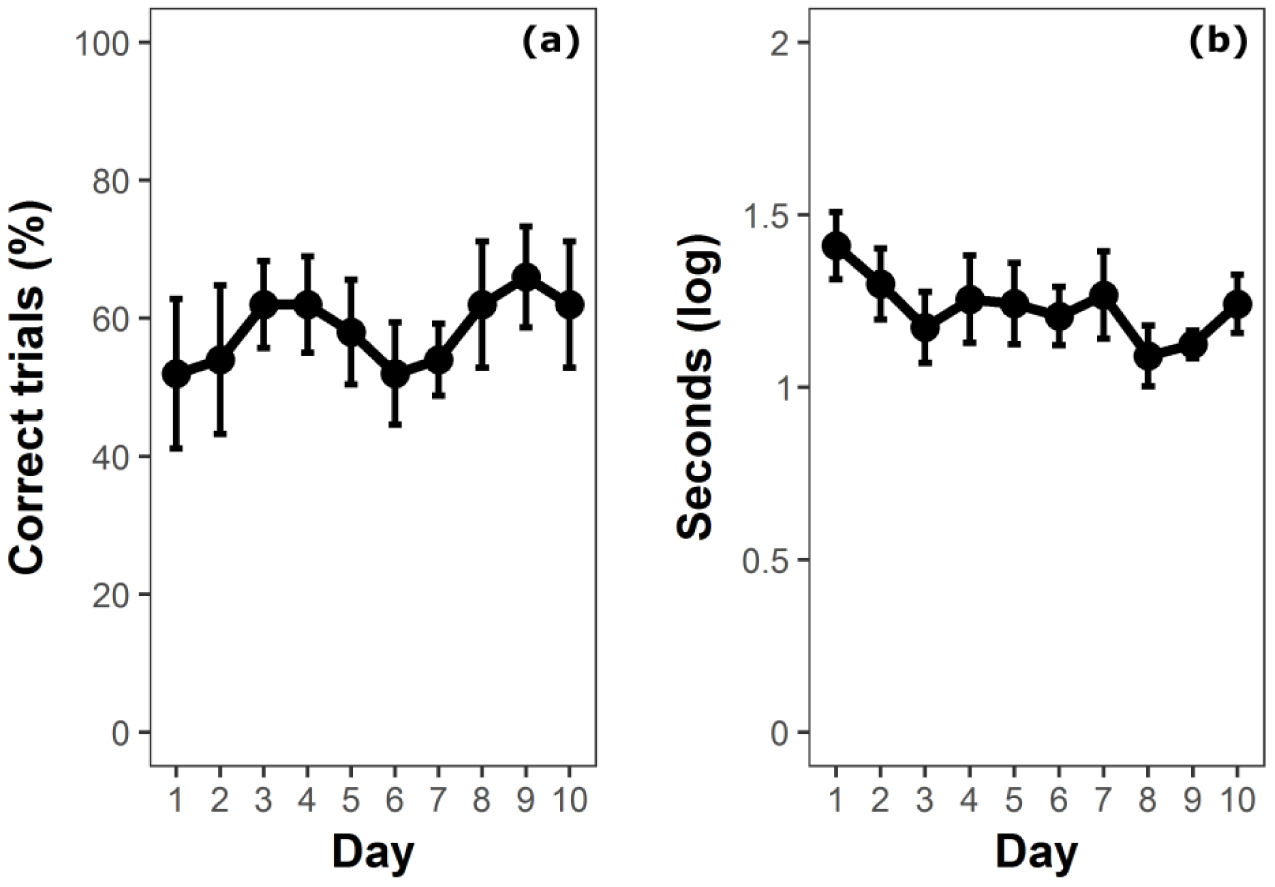
Performance of guppies in the cylinder task (experiment 1). (a) Percentage of correct trials in which guppies did not contact the cylinder (mean ± SEM), and (b) time to solve the task (mean ± SEM) over the 10 days of the test phase.

**Fig. 2.**
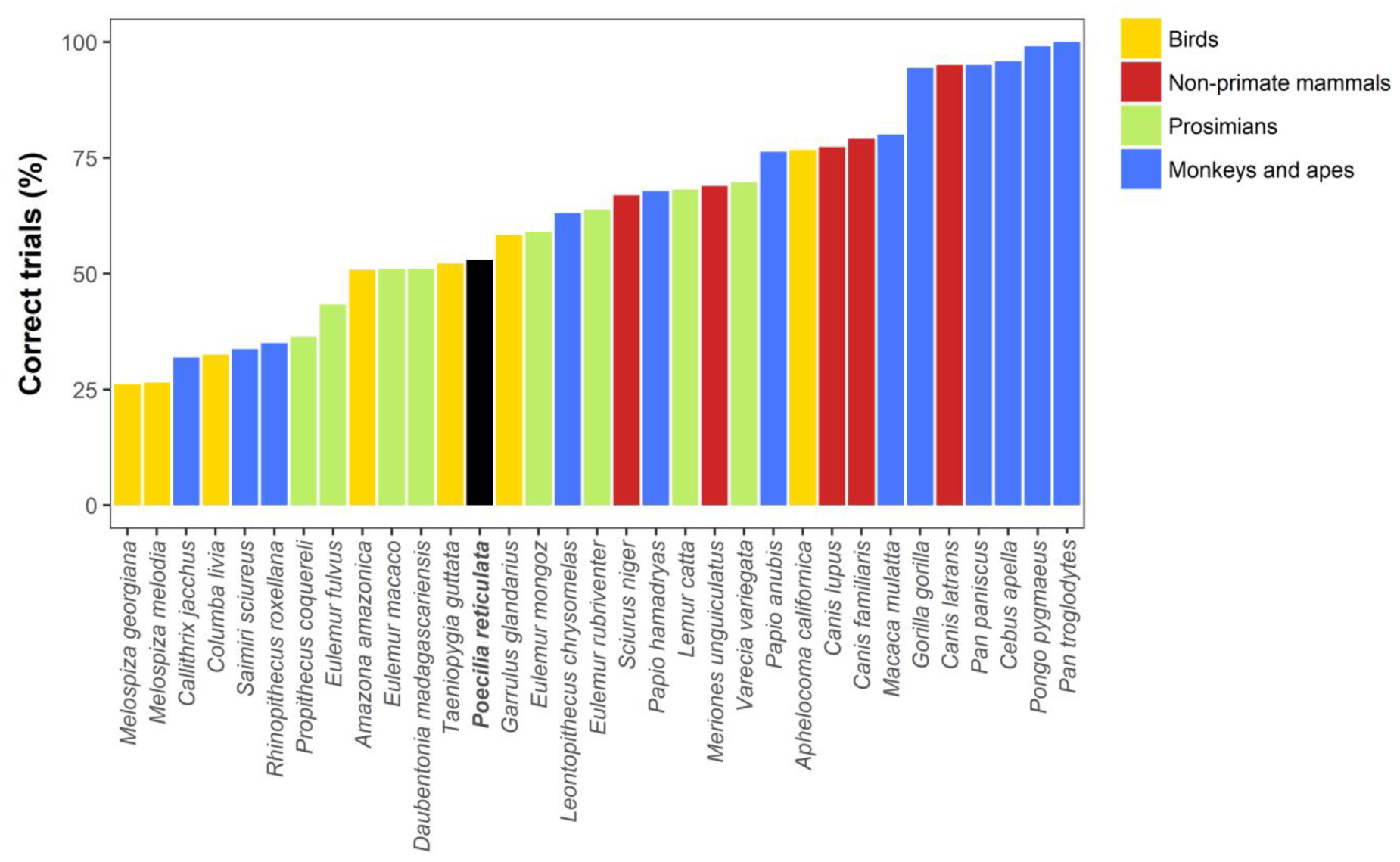
Comparison between the performance of guppies in the cylinder task (black bar) and that of 32 mammalian and avian species tested in the same task by MacLean et al. [20]. Bars represent mean percentage of correct trials. To allow the comparison with the other species, we used the performance of guppies in the initial 10 trials.

### Experiment 2: barrier task

Overall, guppies performed 36.00 ± 21.51 % correct trials in which they reached the stimulus shoal without entering the area delimited by the wings of the transparent barrier. There was clear evidence that the likelihood of a correct trial significantly increased across the trials (GLMM: *χ*^2^_1_ = 24.766, *P* < 0.0001; Fig. 3a), and time spent behind the transparent barrier significantly decreased across the trials (LMM: *χ*^2^_1_ = 31.128, *P* < 0.0001; Fig. 3b). To better compare our experiment with the previous one, we also calculated the guppies’ performance excluding the initial three days, which corresponded to the length of the training phase with the opaque cylinder in experiment 1. When considering only the last two days of training, guppies performed 49.17 ± 34.23 % correct trials.

**Fig. 3.**
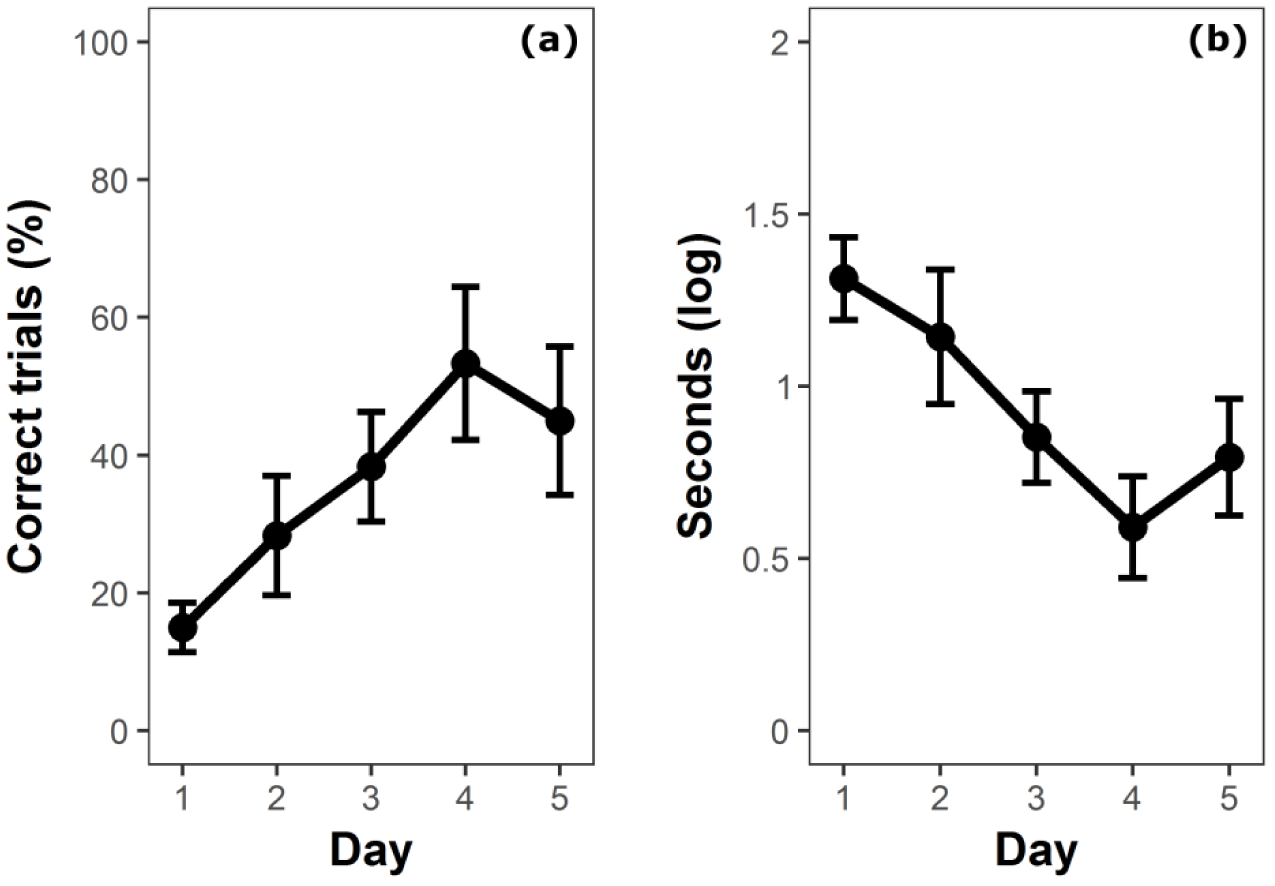
Performance of guppies in the barrier task (experiment 2). (a) Percentage of correct trials in which guppies did not enter the barrier (mean ± SEM) and (b) time to solve the task (mean ± SEM) over the 5 days of the experiment.

## Discussion

In this study, we investigated the ability of a fish species, the guppy, to perform two inhibition motor tasks based on the presence of transparent obstacles between the subject and the goal. Our results indicate that guppies are capable of solving inhibition motor tasks and that their performance is fully comparable to that observed in warm-blooded vertebrates.

In the cylinder task (experiment 1), guppies had to reach a food reward by entering a transparent cylinder from the open lateral sides, rather than trying to approach the food directly. This task was similar to the one adopted in a large study testing 32 species of mammals and birds [20] and thus allowed a direct comparison between guppies and other species. We found no substantial difference between the percentage of correct trials made by guppies, 58 % (53 % if we consider only the first 10 trials), and the average performance of the mammalian and avian species tested by MacLean and colleagues (63 % correct trials; Fig. 2). When apes are not considered, the difference between guppies and warm-blooded species (58 %) is even smaller. Also the individual guppy with lowest performance (38 % correct trials) outperformed many mammalian and avian species.

In contrast with two recent studies on birds [21, 35], we did not find evidence of an increase in the number of correct trials due to training. The absence of change in performance across the 50 test trials also allows to exclude that our measure of inhibitory motor control was affected by the novelty associated with the replacement of the cylinder. Indeed, guppies and other fish species often explore small, armless novel objects introduced in their aquaria [43, 44] and this behaviour might potentially affect performance. It remains to be addressed whether response to novelty might partially explain inter-specific differences in studies with a reduced number of testing trials [20, 21]. The time to solve the task had a small but significant decrease over the trials, which might indicate a small performance improvement due to learning [21]. However, it is likely that, using this protocol, learning mostly occurred in the training phase with the opaque cylinder.

There is evidence that inhibitory control performance might depend on the context and the value of the reward [40, 45]. For example, humans show greater inhibitory control with food than with money reward [40]. Although the procedure of our cylinder task was as close as possible to that adopted for mammals and birds, heterotherms such as guppies may assign a different value to a food reward because of their different metabolic requirements. We controlled for this issue by performing a second experiment in which guppies had to detour a transparent barrier to reach a social reward. The overall performance of guppies in the barrier task tends to be lower than the performance of guppies in the cylinder task. This difference might be explained by the fact that, in the barrier task, during the initial days, the guppies had to learn to detour the barrier and handle the transparency simultaneously; conversely, in the cylinder task, these two phases were separated because the animals were initially trained using an opaque cylinder. Experiments on infants [46], cotton top tamarins [18] and three species of apes [19] showed that subjects initially trained using an opaque barrier performed better than those exposed only to a transparent barrier. In line with this interpretation, at the beginning of the barrier experiment, the performance of guppies was rather poor, less than 20% correct trials. This agrees with a previous study that compared the behaviour of guppies with a transparent and a semi-transparent barrier in 5 test trials, finding reduced detour performance with the transparent barrier [42]. However, in the present study, after three days of experiments (roughly corresponding to the length of the training using the opaque cylinder in experiment 1), the guppies reached a performance very close to that of the cylinder task (50 %). We should additionally consider that the barrier was perhaps more difficult to detour because it was larger than the cylinder and was C-shaped. In the light of these clarifications, the performance of the guppies appears similar for the two different types of reward, and thus this study does not support the hypothesis that the high performance of guppies in experiment 1 was the consequence of a reduced attractiveness of the food reward compared to warm-blooded vertebrates.

The clear increase in the guppies’ percentage of correct trials across testing days in the barrier task was accompanied by a marked decrease in the time taken to solve the task. Both improvements likely indicate that the guppies had learned to handle the transparent barrier. It is interesting to note that a similar improvement was observed in some species (cotton-top tamarins: [18]; orangutans, *Pongo pygmaeus*: [19]) but not in others (gorillas, *Gorilla gorilla*; bonobos, *Pan paniscus*; and chimpanzees, *Pan troglodytes;* [19]). These three latter species performed quite well in the cylinder task [20], and it is still to be addressed whether the differential performance in the two tasks was due to methodological reasons as proposed for guppies.

Only one other study has directly investigated inhibitory performance of fish. Danisman et al. [47] trained cleaner fish, *Labroides dimidiatus*, in a reverse reward contingency task: subjects had to select the smaller food item between two options to receive the larger food item as a reward. They found a poor performance of cleaner fish with none of the eight subjects being able to learn the task. Many other species did not succeed in learning to solve this task (e.g., chimpanzees: [48]; Japanese macaques, *Macaca fuscata*: [49]; cotton-top tamarins: [9]; black and brown lemurs, *Eulemur fulvus* and *E. macaco*: [17]). The difference between the study on cleaner fish and our study on guppies is likely due to the large difficulty of the reverse reward contingency task. To address this point, we need to gather more data on the performance of fish in other inhibitory control tasks. Among the others, it will be important to focus on tasks requiring self-control (i.e., the choice between alternatives with different values and different costs), because self-control is generally considered the most challenging aspect of inhibitory control [36].

An efficient inhibitory control has been usually considered typical of humans and primates [11, 12], and it has been shown to positively correlate with brain size in a recent comparative study [20]. As guppy’s brain is more than 100 times smaller than the brain of the smallest species included in that comparative study, the performance of the guppies in the cylinder task is exceedingly higher than would be expected based on brain size. Together with other evidence [21], this suggests that brain size alone cannot explain the large differences in inhibitory motor control observed among species.

In MacLean’s study, the main predictor of inhibitory motor control performance was absolute brain size [20]. Perhaps this relationship only holds considering a sample of species with a limited range of body size or within a restricted taxonomic group. As recently discussed by Herculano-Houzel [50], brain mass is only a proxy for the neuronal capability devoted to complex information processing. If larger bodies require larger brains to operate, then in larger species only part of the increase in brain mass can contribute to behavioural complexity. However, controlling for the whole brain allometry is unlikely to account for the performance of guppies, as fish have, on average, a relative brain weight ten times smaller than mammals and birds [51].

Another important issue to be considered is that the brain of different species can differ in structure at different scale levels, and these differences are expected to increase with increased phylogenetic distance. For example, neural density is extremely variable both within mammals and between mammals and birds [50]. Recently, Kabadayi et al. [21] found that in three corvid species, performance in a inhibitory motor control task was much higher than the average performance found in mammals, although their brain mass is much smaller. This result could partly be explained by the fact that the forebrain of several bird species contains many more neurons compared to that of mammals [52]. As another example, some insects are capable of exceptional cognitive performance despite having a brain that is extremely small even compared to a small fish like the guppy; it was suggested that this may be related to characteristics of the neural circuits that present wide differences between vertebrates and arthropods [53].

Though they belong to the same clade, the modern ray-finned fishes (to which teleosts belong) diverged approximately 450 million years ago from the line of fish that gave origin to land vertebrates. In addition, a major genomic rearrangement – whole-genome duplication – occurred in the line leading to teleosts soon after the separation; there is now evidence that this event produced a significant enrichment of the set of genes available for the evolution of novelties in the nervous system of this vertebrate group [54]. Therefore, the brains of teleost fish and land vertebrates evolved in large part independently and may show a very different anatomical and cytoarchitectonic structure. However, the cytoarchitectonic structure of the teleost brain and the localisation of the functions studied here are less well known compared to warm-blooded vertebrates and therefore any conclusion on this topic is premature.

A second important factor that can explain interspecific differences in cognitive abilities across all vertebrates is the selective pressure exerted by the environment in which a species evolved. Several ecological factors have been suggested to promote the evolution of inhibitory control. For example, species that typically feed on moving prey might show more impulsiveness [11]. Alternatively, species with a complex social environment may have been selected for greater inhibitory control, a hypothesis that has found some support [55]. Another possibility is that inhibitory control evolves as a by-product of selection on other behaviours and cognitive functions. As the capacity to inhibit prepotent but unfavourable responses is an important prerequisite for a wide range of cognitive tasks [6, 7, 8, 9, 10], it is conceivable that selection acting on these cognitive functions can indirectly select for high inhibitory control.

## Materials and methods

### Ethical statement

The experiments adhere to the current legislation of our country (Decreto Legislativo 4 marzo 2014, n. 26) and were approved by the Ethical Committee of Università di Padova (protocol n. 33/2015).

### Subjects

The subjects were adult female guppies of an ornamental strain (“snakeskin cobra green”) bred in our laboratory since 2012. We tested 10 guppies in experiment 1 and 12 guppies in experiment 2. The maintenance tanks (400 L), which housed guppies before the experiments, had a gravel bottom, abundant natural and artificial plants, water filters, and 15-W fluorescent lamps (12h:12h light/dark photoperiod). We kept water temperature at 26 ± 1 °C and fed the fish with commercial food flakes (Fioccomix, Super Hi Group, Ovada, Italy) and *Artemia salina* nauplii three times per day.

### Experiment 1: cylinder task

#### Apparatus

In an 80 × 40 × 38 cm tank filled with 30 cm of water, we built a green plastic apparatus in the shape of an hourglass (Fig. 4a) similar to the ones adopted in previous studies on guppies [33, 56, 57]. The central corridor (10 × 10 cm) connected two test compartments (28 × 40 cm) by means of two 10 × 8 cm guillotine transparent doors. Each trapezoidal compartment beside the corridor had green net walls and housed one immature guppy as a social companion, abundant vegetation, and a water filter. One 18-W fluorescent lamp and one video camera were placed above each test compartment.

We used two types of cylinders of the same size (4 cm length, 3.5 cm diameter). During the training phase, the cylinder was opaque (green plastic), whereas during the test phase, the cylinder was transparent (an acetate sheet). Both cylinders were glued above a green plastic sheet (4 × 4 cm).

**Fig. 4.**
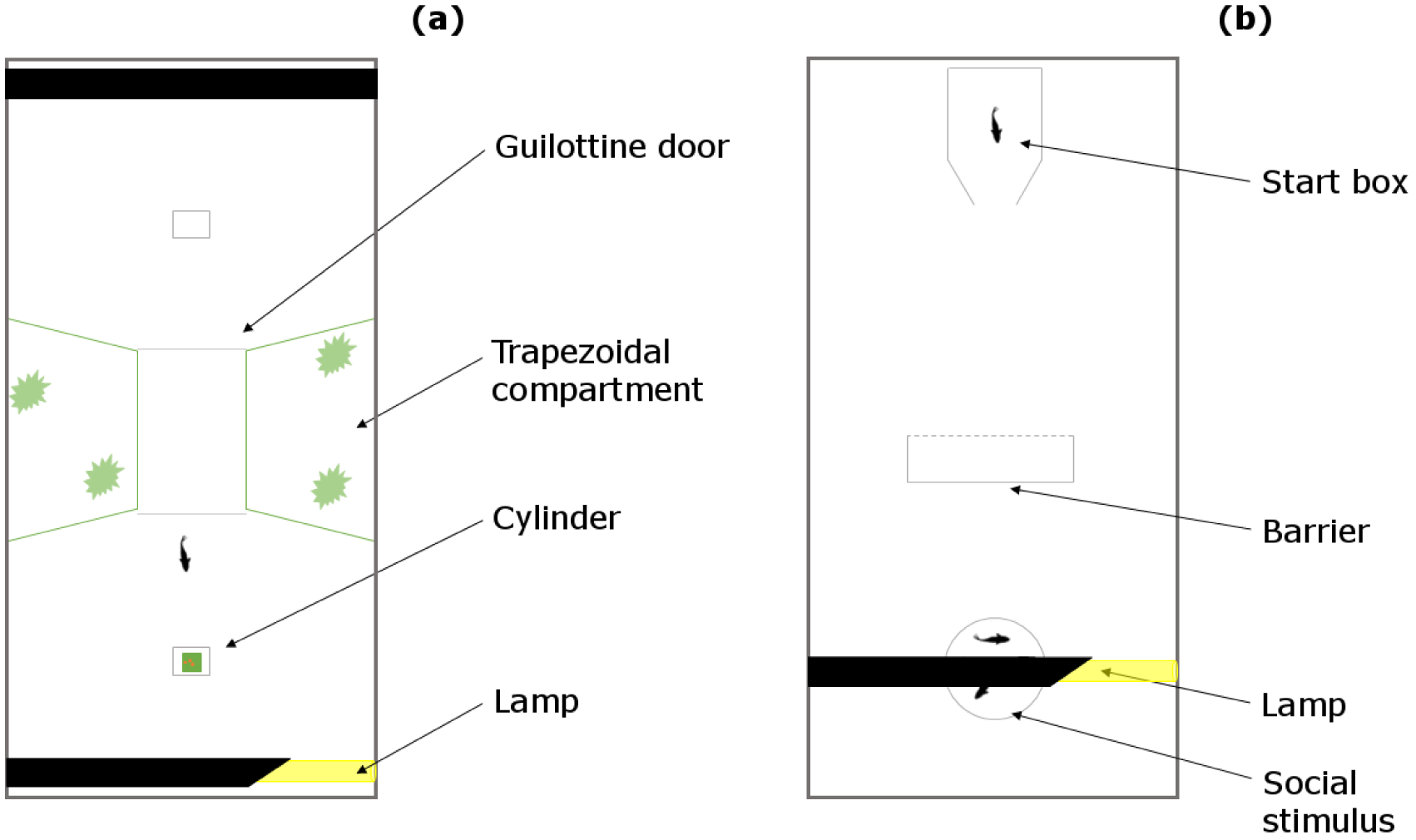
View from above of the apparatuses used in (a) experiment 1, and in (b) experiment 2.

#### Familiarization with the apparatus and the procedure

Three days before the beginning of training, we randomly selected a female in the maintenance tanks and moved it to the apparatus, together with one adult male and three juveniles to avoid social isolation. We fed the subject five times each day in the two test compartments alternately, in order to simulate the procedure of the following days (see below). Before feeding the subject, we moved the male companion to a 60 × 40 × 38 cm tank with immature guppies as social companions, vegetation, and water filters, and we fed it thereby. Then, we inserted a Pasteur pipette with crumbled flakes mixed with water into the test compartment opposite the one with the subject. We slowly moved the pipette to attract the subject; when the subject entered this test compartment, we closed the guillotine door and slowly released the food from the pipette. This was done to train the fish to get the food from the pipette. Five guppies that did not learn to feed from the pipette were discarded and substituted with new subjects. The day before the experiment started, the immature companions were removed from the tank, with the exception of the two in the trapezoidal compartments.

#### Training phase

The subject was trained to feed inside the opaque cylinder. We performed 5 trials per day, in which the position of the opaque cylinder was alternated between the two test compartments. Thirty minutes before the experiment started, the male companion was removed from the tank, and the female was confined in one test compartment. The experimenter placed the cylinder into the other test compartment at a distance of 15 cm from the guillotine door. Then, the experimenter inserted the Pasteur pipette and showed it to the subject confined behind the guillotine door. After ensuring that the subject was looking in the direction of the pipette, the experimenter inserted food inside the cylinder and opened the guillotine door. The subject had 30 min to find the food; after this period the trial was considered null and was repeated thereafter. If the subject entered the cylinder, we waited 15 min and then started the next trial in the opposite test compartment. Based on the video recordings, we measured whether the first attempt to reach the food was through the front of the cylinder (incorrect trial) or from the open sides (correct trial). Subjects had to achieve four out of five correct trials in a day to pass to the test phase.

#### Test phase

In the test phase, we used the transparent cylinder but other details were identical to the training phase. Based on the video recordings, we scored whether the subject first attempted to retrieve food through the cylinder (incorrect) or from the side (correct) and the time to enter the cylinder in each trial. Since the procedure of the cylinder task has never been used in fish, in a pilot experiment we analysed the reliability of our measures of performance. We performed 40 trials in which the performance of 6 guppies was scored live by the experimenter and from the video recordings by two other scorers. Regarding correct versus incorrect trials, the live score and one score from the recordings were identical in all the trials, whereas the second score from the recordings differed in 1 out of 40 trials (2.5 % of the trials). Regarding the time to solve the task, the three scores were highly correlated (Spearman’s rank correlation: *ρ* = 0.997, *P* < 0.0001; *ρ* = 0.980, *P* < 0.0001; *ρ* = 0.981, *P* < 0.0001). These analyses revealed that our measures of performance were robust and repeatable. Subjects were tested for 10 days (5 trials each day, 50 total trials), the only difference from the original method of MacLean and colleagues [20] which performed 10 test trials.

### Experiment 2: barrier task

#### Apparatus

The apparatus was an 80 × 40 × 36 cm tank covered with a white plastic sheet and filled with 10 cm of water (Fig. 4b). In one of the short sides of the tank, we built a white start box (15 × 10 × 20 cm). The social stimulus was a shoal of 4 female guppies confined in a transparent sector of the apparatus (11 cm diameter, 18 cm height). We positioned the transparent barrier (18 × 18 cm), C-shaped by means of two white plastic wings (18 × 5 cm), at a distance of 30 cm from the start box and 15 cm from the stimulus. An 18-W fluorescent lamp placed above the stimuli illuminated the apparatus and a video camera recorded the trials.

#### Procedure

One week before the beginning of the experiment, we moved each individual subject from the maintenance tank to a 50 × 20 × 38 cm ‘home tank’ with immature guppies as social companions, vegetation, and a water filter. The experiment consisted of a series of 25 test trials subdivided over 5 days (5 trials per day). The length of the experiment was reduced compared to experiment 1 for ethical reasons because this procedure was presumably more stressful for the subjects. We placed the stimuli in the sector of the apparatus 30 min before the first trial. Successively, one subject was netted from its home tank, transported in a plastic jar and gently inserted into the start box, from which it could swim to the stimulus. From the video recordings, we scored whether the subject reached the stimulus shoal by entering into the area delimited by the wings of the barrier (incorrect trial) or not (correct trial) and the time spent within this area. After the subject joined the shoal, we left it undisturbed for 5 min as a reward before starting the following trial. Four subjects that did not attempt to join the stimulus shoal within 20 min were substituted. At the end of the 5 daily trials, the subject was moved to the home tank.

As in the cylinder task, we analysed the reliability of our measures of performance. A second experimenter re-analysed the video recordings of 40 trials by 8 guppies. The binary measure of performance, correct versus incorrect trials, differed between the two scores in 1 out of 40 trials (2.5 % of the trials). The time to solve the task was highly correlated between the two scores (Spearman’s rank correlation: *ρ* = 0.987, *P* < 0.0001).

#### Statistical analysis

Analyses were performed in R version 3.2.2 (The R Foundation for Statistical Computing, Vienna, Austria, http://www.r-project.org). For both experiments, we analysed the outcome of the trials (correct or incorrect) with generalized linear mixed-effects models for binomial response distributions (GLMMs, ‘glmer’ function of the ‘lme4’ R package) fitted with the trial number as covariate (to examine whether performance improved over trials) and individual ID as random effect. To compare the average score of guppies in the cylinder task with the data set of 32 mammalian and avian species from MacLean et al. [20], we computed the percentage of correct trials in the first 10 trials. In experiment 1, we also compared the proportion of correct trials in the last day of the training phase of each subject versus the test phase using paired-sample *t* test. We analysed time performance (time to reach the reward in experiment 1 and time spent trying to pass thought the barrier in experiment 2) using linear mixed-effects models (LMMs, ‘lmer’ function of the ‘lme4’ R package) fitted with the trial as covariate, after log transformation due to a right-skewed distribution, and individual ID as random effect.

## Acknowledgements

The authors would like to thank Alice Andreoli, Martina Zanutto, and Pietro Maconi for their help in testing the animals. This research was supported by DOR Grant to A.B. from Università di Padova.

## Author contributions

T.L.-X., E.G., and A.B. developed the study concept and design. E.G. collected the data. T.L.-X. analysed the data and drafted the manuscript. All authors provided critical revisions and approved the final version of the manuscript.

